# Brightest path tracing: A Python package to trace the brightest path in 2D and 3D images

**DOI:** 10.1101/2023.07.16.549233

**Authors:** Vasudha Jha, Robert H. Cudmore

## Abstract

Brightest path tracing is a widely used image processing technique in several fields including biology, geography, and geology. However, despite the availability of many image processing libraries in Python, few offer an out-of-the-box implementation of a brightest path tracing algorithm. This paper presents a Python package, `brightest-path-lib`, that efficiently finds the path with maximum brightness between points in a 2D or 3D image. An example graphical user interface is provided as a Napari plugin. Taken together, the package and plugin provide a powerful and extensible tool for users to efficiently trace structures of interest in 2D or 3D images, regardless of the type of structure being analyzed.

## Introduction

Brightest path tracing is an image processing technique which involves identifying the brightest points in an image and then connecting them to form a path. This path can provide valuable information about the structure and organization of the object being traced, such as their connectivity and morphology. Brightest path tracing is commonly used in neuroscience and medical imaging to trace structures such as neurons or blood vessels, aiding in disease diagnosis and treatment.

Python has a thriving ecosystem of image processing libraries that provide a broad range of functionalities, from basic operations like image manipulation and saving to advanced tasks like object detection and machine learning-based image analysis. Some of the widely used image processing libraries in Python include OpenCV (Bradski 2000), scikit-image (Walt et al. 2014), imageio (Klein et al. 2023), and Pygame (Pygame 2023). Although these libraries support implementing search algorithms such as A* (Hart, Nilsson, and Raphael 1968) and Dijkstra’s algorithm (Dijkstra 1959) to find the shortest path between two points in an image, they lack built-in support for brightest path tracing. While other software tools like ImageJ (Abràmoff, Magalhães, and Ram 2004) offer solutions for neuronal tracing, such as the Simple Neurite Tracer (Longair, Baker, and Armstrong 2011; Arshadi et al. 2021), they are implemented in Java and may pose challenges when integrating with Python-based workflows. Here, we present the `brightest-path-lib`, a Python package designed to fill this gap by providing native Python implementations of informed search algorithms tailored specifically for brightest path tracing, making it easier to integrate into existing Python-based image processing pipelines.

## Results

The `brightest-path-lib` package is available on PyPi for installation using pip and the source code is provided on GitHub (https://github.com/mapmanager/brightest-path-lib). The package includes efficient algorithms to trace the path between points with the maximum brightness in 2D and 3D images (Figure 1). Two search algorithms are provided including A* Search and Bidirectional Search (Pijls and Post 2009; Wink, Niessen, and Viergever 2000).

**Figure 1:**
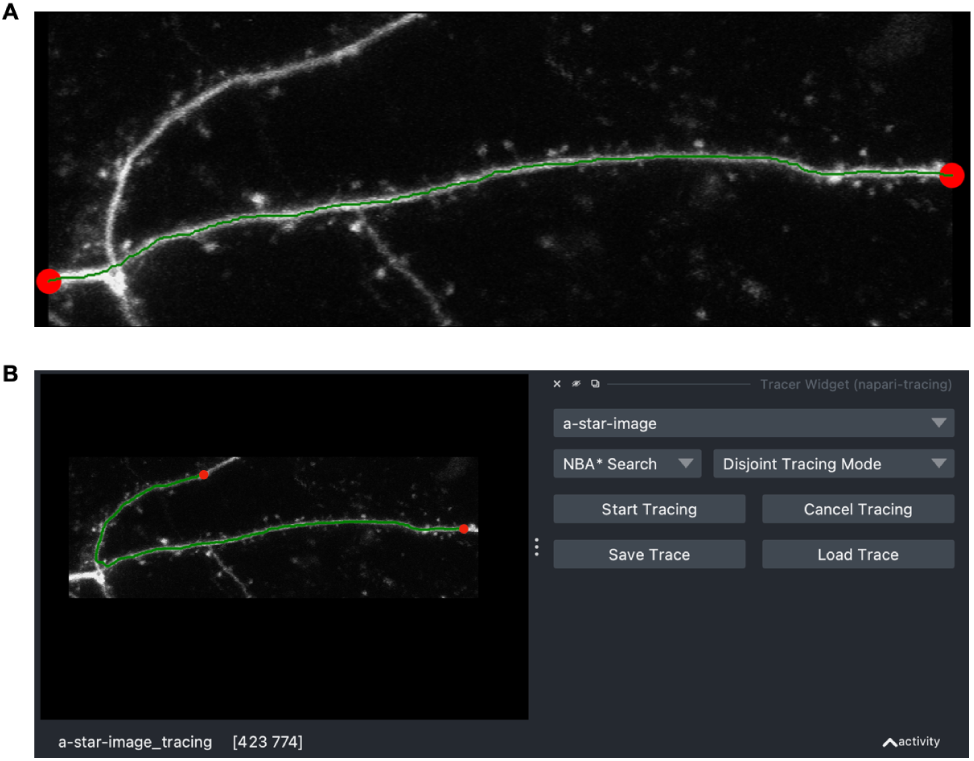
Example results of brightest-path tracing of a neuronal dendrite. (A) An example tracing performed from a Python script. The start and stop points are red circles, the final brightest path is in green. The grayscale image is a maximal intensity projection of a 3D image volume. (B) An example screenshot of the Napari Tracing Plugin. On the left is the image, start and stop points (red), and the final brightest path (green). On the right is the user interface for tracing including controls to select the tracing algorithm, start and cancel the tracing, and finally to save and load the tracing as an SWC file.

Detailed application programming interface (API) and user documentation as well as example work-flows are provided, enabling users to quickly get started with the package and understand its functionalities. Finally, the use of open-source licensing, and Python’s extensive usage in the scientific community ensures a vast community of users and developers who can contribute to the package ‘s continued growth and improvement.

To allow non-programming users to utilize the `brightest-path-lib` package, we have developed a Napari (Napari 2023) plugin called `Napari Tracer Plugin` (available at https://github.com/mapmanager/napari-tracing) that integrates the `brightest-path-lib` to provide a graphical user in-terface (GUI) for real-time visualization of the path and the image, making it easier to understand the object’s structure and organization being traced (Figure 1). Finally, tracings are saved in the SWC neuron morphology file format (Stockley et al. 1993).

## Discussion

Together, the `brightest-path-lib` package and the Napari plugin provide a powerful set of tools for both programmers and end-users to efficiently trace structures of interest. Moreover, the plugin is designed to work seamlessly with other Napari plugins, allowing for more integrated and streamlined workflows. This feature further extends the capabilities of the `brightest-path-lib` and `Napari Tracer Plugin` beyond their core functionalities, providing users with a more comprehensive suite of tools for image analysis and visualization.

## Acknowledgements

VJ and RHC were supported by an NIH grant (1RF1MH123206-01A1) and a Chan Zuckerberg Initiative grant (2022-252611).

